# CFSA: Comparative Flux Sampling Analysis as a Guide for Strain Design

**DOI:** 10.1101/2023.06.15.545085

**Authors:** R.P. van Rosmalen, S. Moreno-Paz, Z.E. Duman-Özdamar, M. Suarez-Diez

## Abstract

Genome-scale metabolic models of microbial metabolism have extensively been used to guide the design of microbial cell factories, still, many of the available strain design algorithms often fail to produce a reduced list of targets for improved performance that can be implemented and validated in a step-wise manner. We present Comparative Flux Sampling Analysis (CFSA), a strain design method based on the extensive comparison of complete metabolic spaces corresponding to maximal or near-maximal growth and production phenotypes. The comparison is complemented by statistical analysis to identify reactions with altered flux that are suggested as targets for genetic interventions including up-regulations, down-regulations and gene deletions. We applied CFSA to the production of lipids by *Cutaneotrichosporon oleaginosus* and naringenin by *Saccharomyces cerevisiae* identifying engineering targets in agreement with previous studies as well as new interventions. CFSA is an easy-to-use, robust method that suggests potential metabolic engineering targets for growth-uncoupled production that can be applied to the design of microbial cell factories.

## 1 Introduction

Microbial cell factories (MCF) are microorganisms engineered for the production of bio-molecules that thrive on renewable carbon sources. A broad range of bio-molecules can be produced using MCF from non-edible feedstocks such as recalcitrant biomass or industrial waste streams thereby providing sustainable replacements for production systems based on fossil fuels [1]. MCF design requires the choice of an appropriate host strain (chassis), and selection of a suitable available pathway, the discovery of new pathways, or the design of synthetic pathways for new-to-nature compounds. Still, industrial feasibility requires extensive engineering to improve MCF performance [2].

GEnome-scale Metabolic models (GEM) comprehensively represent cellular metabolism and allow the simulation of metabolic fluxes and the prediction of cellular phenotypes. They have largely been used to identify metabolic engineering targets to optimize pathway performance [3]. Machado et al. [4] classify strain design algorithms relying on GEMs in two main groups: methods based on the analysis of elementary flux modes (EFM), and those based on the optimization of an objective function.

EFMs are minimal sets of reactions that can jointly operate at steady-state such that all steady-state solutions can be described as a combination of EFMs [5]. They provide an unbiased framework to explore the metabolic space but have limited scaling potential and applicability to larger models. Instead, related approaches, based on minimal cut sets (MCS) such as minimal metabolic functionality (MMF) and FluxDesign, scale better to genome-scale [4, 6].

Optimization-based approaches rely on the simulation of metabolic fluxes of *wild type* and/or mutant strains using an objective function. Many of these methods use Flux Balance Analysis (FBA) to calculate fluxes and therefore only explore one of the multiple flux distributions that can lead to the optimal objective ignoring the rest. In this way, OptKnock [7] and derived algorithms such as RobustKnock [8], OptGene [9], and OptCouple [10], aim at identifying gene knock-outs to couple the production of the compound of interest to the production of biomass using growth as the objective function. Other methods such as OptForce [11] and OptDesign [12] are used to predict modulation of gene expression, including up and down-regulations. They compare simulated fluxes of growth and production phenotypes and minimize the number of interventions required for the overproduction phenotype. Although flux variability analysis (FVA) might be used to explore feasible flux ranges and constrain the solution space, comparisons among phenotypes are based on non-unique FBA solutions. Alternatively, flux sampling allows the exploration of the full space of feasible flux distribution of each reaction given a set of constraints on the metabolic model [13]. Contrarily to FBA, flux sampling does not require the selection of an objective function, which biases FBA predictions. Advantages of flux sampling have been exploited to evaluate metabolic flux differences between different conditions [14], but have not yet been expanded to strain design.

Described strain design algorithms focus on the identification of growth-coupling strategies, where production becomes a requirement for growth [7–10]. This strategy is suitable for experimental implementation and further optimization using adaptive laboratory evolution but requires multiple simultaneous interventions. Alternatively, strains with growth-uncoupled production can be used in two-stage fermentation processes where the growth and production phases are sequential. This strategy can alleviate metabolic stress and improve productivity [15, 16].

We present Comparative Flux Space Analysis (CFSA), a model-guided strain design approach based on extensive sampling of the feasible solution space in alternative scenarios. Growth and production phenotypes are simulated and compared, also with a growth-limited scenario, which serves as a negative control for down-regulation targets. Flux distributions are statistically compared resulting on the identification of potential down-regulations, knock-outs and over-expressions targets leading to growth-uncoupled increased production. As a proof of concept, we use CFSA to identify metabolic engineering targets for lipid production by *Cutaneotrichosporon oleaginosus* and naringenin production by *Saccharomyces cerevisiae* and compare them with available data.

## 2 Methods

### 2.1 Comparative Flux Sampling Analysis (CFSA)

To implement CFSA a GEM of the desired organism including the production pathway of choice is required. As the first step, media conditions (e.g. substrate uptake rates, aerobic/anaerobic growth…) are specified. Model reactions are grouped in seven categories to facilitate later filtering: *required* reactions including the growth and maintenance reactions; *not biological* reactions including boundary, exchange, sink and demand reactions; *blocked* reactions (unable to carry flux); reactions *without associated genes*; *essential* reactions; reactions *containing essential genes* and *transport* reactions.

Reaction fluxes are sampled from the metabolic solution space in three scenarios: growth, slow growth, and production. In the growth and production scenarios, the optimality parameter ensures that sampled flux distributions result in at least a specified fraction of the optimal growth or production predicted by FBA by constraining the lower bound of the biomass or product exchange reactions. In the slow growth scenario, the maximum growth rate compatible to the specified minimal production rate is calculated and used as an upper bound for the biomass synthesis reaction. To limit the solution space a parsimonious FBA approach [17] is implemented by introducing an additional constraint to limit the total sum of fluxes to the minimum value compatible with maximal growth given by the flux fraction parameter. This extra constraint is applied in the production and slow growth scenario simulations to limit unrealistic futile cycles.

The Optimal Gaussian Process (OptGP) sampler [18] implemented in cobrapy [19] is used to model the distribution of the target space and iteratively sample from this distribution. A thinning parameter is used to reduce the correlation between samples. Invalid samples (i.e. those that do not meet constraints specific to each scenario) are discarded. Last, the Geweke diagnostic is used to calculate the chain convergence for each process, and samples corresponding to reactions whose distributions have not converged are discarded [20].

For each reaction, the two-sample Kolmogorov-Smirnov (KS) test is used to compare samples from different scenarios and determine if they belong to the same continuous distribution. Potential targets are selected if, for a specific reaction, distributions differ in the scenarios based on KS-statistics and p-values corrected for multiple testing using Bonferroni. The p-value and KS cutoffs can be adjusted by the user. Besides, reactions whose fluxes correlate with fluxes through the biomass synthesis reaction, or that are not associated to genes (through a gene-protein-reaction association) are discarded as potential targets. Only reactions whose absolute change in flux between the growth and production scenario is bigger than a user-specified threshold are considered to be suitable targets. Similarly, targets can be filtered based on the standard deviation of the samples taken in the production scenario.

Potential targets are then divided into over-expression or down-regulation targets depending on whether the mean fold change comparing growth and production scenarios is above or below 1. Knock-down targets that correspond to non-essential genes are classified as possible knock-out targets. Reactions are clustered based on the correlation of the absolute fluxes between samples in order to identify redundant targets (i.e. belonging to the same metabolic pathway).

### 2.1 Selected applications

#### 2.2.1 Production of lipids by *C. oleaginosus*

The *i* NP636 *Coleaginosus* ATCC20509 genome-scale metabolic model was manually curated to provide all fatty acid elongation reactions [21]. In total 7 reactions were added, 1 of them in the cytoplasm and 6 of them in the mitochondria, and all reactions were associated with the corresponding genes. Details of the model curation can be found on [GitLab](https://gitlab.com/wurssb/Modelling/sampling-tools/-/blob/master/example_filtering_yeast.ipynb?ref_type=heads). Glycerol and urea were used as the carbon and nitrogen sources respectively and nitrogen-depleted biomass composition was assumed. CFSA was used (optimality = 0.90, flux fraction = 1.25, KS1 = KS2 ≥ 0.75, mean absolute change ≥ 0.01, standard deviation in production ≤ 50) using the lipid synthesis reaction (lipid_synthesis) as a target for the production scenario. The feasibility of selected targets was evaluated based on previous studies.

#### 2.2.2 Production of naringenin by *S. cerevisiae*

The Yeast8 genome-scale metabolic model of *S. cerevisiae* [22] was modified to include the naringenin production pathway (Table 1) and glucose was the selected carbon source. CFSA was used (optimality = 0.90, flux fraction = 1.25, KS1 = KS2 *≥* 0.75, mean absolute change *≥*0.01, standard deviation in production *≤* 50) with the naringenin exchange reaction (EX_NAR) as an objective for the production scenario. Two proteomic datasets representative of *S. cerevisiae* aerobic growth on glucose during shake flask and chemostat fermentations were used as additional filters so only detected proteins were consider suitable down-regulation targets [23, 24].

**Table 1:**
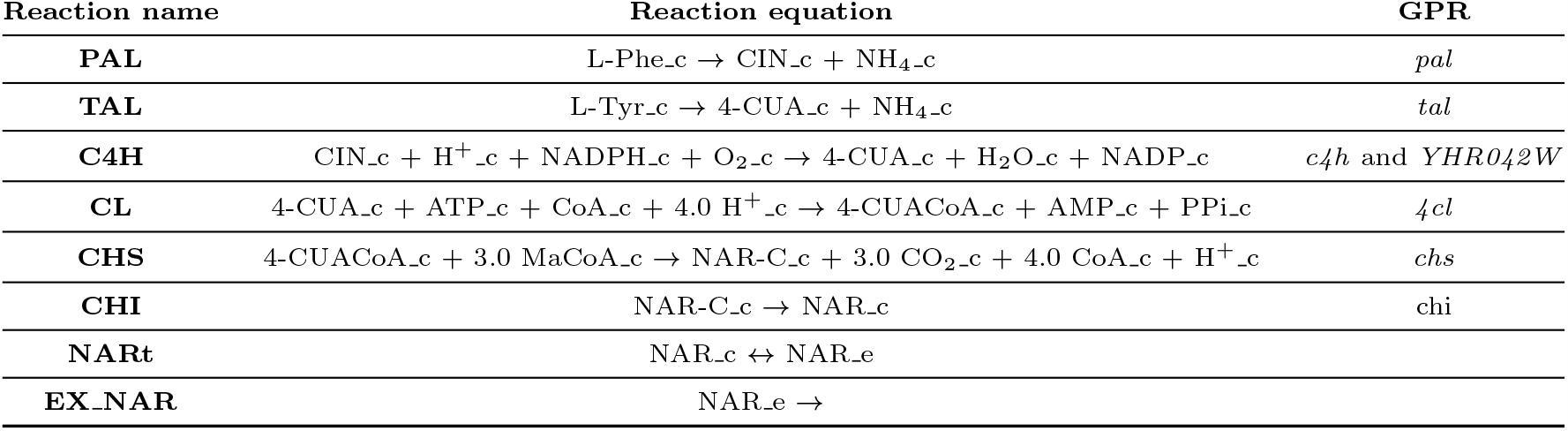
Naringenin pathway reactions added to Yeast8. Reaction names in the model, reaction equations and Gene-Protein-Reaction (GPR) rules are included. PAL, phenylalanine ammonialyase; TAL, tyrosine ammonia-lyase; C4H, cinnamate 4-hydroxylase; CL, 4-coumarate-CoA ligase; CHS, naringenin-chalcone synthase; CHI, chalcone isomerase; NARt, naringenin transport; EX NAR, naringenin exchange; L-Phe, L-phenylalanine; CIN, cinnamate; L-Tyr, L-tyrosine; 4-CUA, 4-coumarate; CoA, Coenzyme-A; CUACoA, 4-coumaroyl-CoA; MaCoA, malonyl-CoA; NAR-C, naringenin chalcone; NAR, naringenin. _c and _e designate metabolites in the cytoplasm and the extracellular space respectively.

| Reaction name | Reaction equation | GPR |
| --- | --- | --- |
| **PAL** | $L\text{-Phe}_c \rightarrow CIN_c + NH_4_c$ | *pal* |
| **TAL** | $L\text{-Tyr}_c \rightarrow 4\text{-CUA}_c + NH_4_c$ | *tal* |
| **C4H** | $CIN_c + H^+_c + NADPH_c + O_{2,c} \rightarrow 4\text{-CUA}_c + H_2O_c + NADP_c$ | *c4h* and *YHR042W* |
| **CL** | $4\text{-CUA}_c + ATP_c + CoA_c + 4.0 H^+_c \rightarrow 4\text{-CUACoA}_c + AMP_c + PPi_c$ | *4cl* |
| **CHS** | $4\text{-CUACoA}_c + 3.0 MaCoA_c \rightarrow NAR\text{-}C_c + 3.0 CO_{2,c} + 4.0 CoA_c + H^+_c$ | *chs* |
| **CHI** | $NAR\text{-}C_c \rightarrow NAR_c$ | *chi* |
| **NARt** | $NAR_c \leftrightarrow NAR_e$ | |
| **EX_NAR** | $NAR_e \rightarrow$ | |

## 3. Results

### 3.1 Comparative Flux Sampling Analysis (CFSA)

CFSA was implemented in python v.3.7 using the cobrapy v 0.26.2 toolbox and is available on [GitLab](https://gitlab.com/wurssb/Modelling/sampling-tools/-/blob/master/example_filtering_yeast.ipynb?ref_type=heads). Statistical analysis methods based on the Kolmogorov-Smirnov test were implemented using scipy stats package. The CFSA output consists of a first excel file with sampling results for all model reactions that is used as input for filtering. After filtering (based on user defined parameters), a new excel file containing the filtered results is generated. This file contains the suggested reaction targets and their associated genes as well as possible off-targets caused by multifunctional enzymes. It provides a summary of the sampling results including the mean fluxes in the growth and production scenario as well as the absolute flux change and the reaction equation. The complete files for both case studies are available in [GitLab](https://gitlab.com/wurssb/Modelling/sampling-tools/-/blob/master/example_filtering_yeast.ipynb?ref_type=heads). Additionally, distribution graphs for suggested targets are generated and can be checked before experimental implementation.

#### 3.1.1 Distribution graphs

A distribution graph can be generated for each reaction in the model. It shows the distribution of sampled fluxes in each scenario: growth, production, and slow growth (*i*.*e*. in how many samples a reaction had a specific flux). Genes are classified as over-expression targets when the absolute mean flux through the corresponding reaction is higher in the production scenario compared to the growth scenario and the flux distributions do not overlap (Figure 1, top left). Similarly, genes are classified as down-regulation targets when the absolute flux through the corresponding reactions is lower in the production scenario compared to the growth scenario and distributions don’t overlap (Fig 1, top right). The most extreme case of a down-regulation, a knock-out, is obtained when, for a down-regulation target, the flux in the production scenario is zero and the gene is not classified as essential (Fig 1, bottom left).

**Figure 1.**
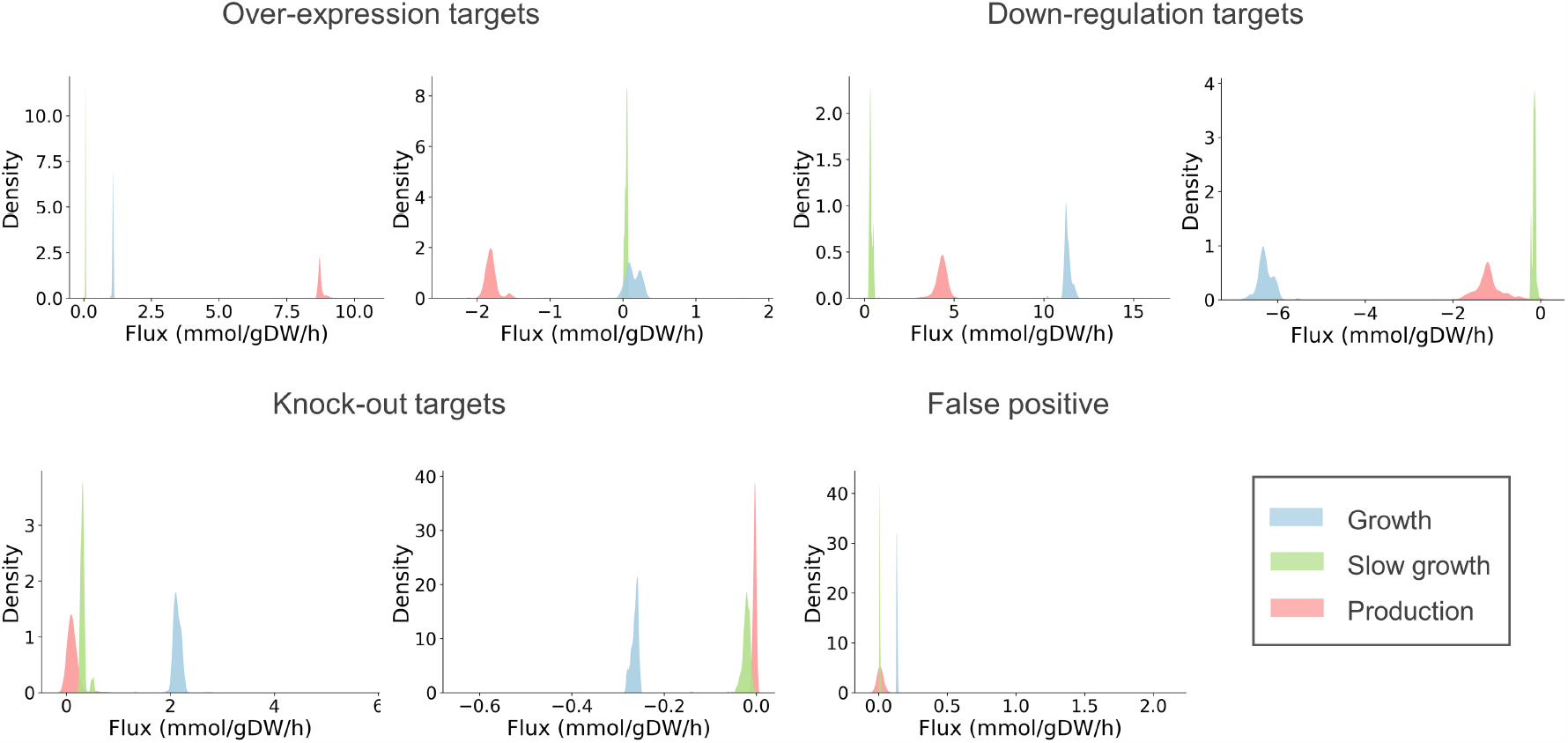
Example of flux distribution plots illustrating behaviors of candidate targets. In each panel, the x-axis represents possible flux values, and the y-axis represents the frequency of each flux value obtained when sampling the solution space, normalized to an area of 1.

The slow growth scenario is used to reduce the number of false positive down-regulation targets. Growth and production are competing objectives and often low fluxes obtained in the production scenario are not related to increased production but to a decreased growth rate (e.g. fluxes through reactions for biomass components). In order to avoid the identification of these genes as down-regulation targets, reactions where the production and slow growth distributions overlap are considered false positives (Fig 1, bottom right).

Distribution graphs also show the allowed variability of a reaction flux. Reactions with sharp distributions require a specific flux to obtain high production. Conversely, reactions with broad distributions are allowed to carry different fluxes without impacting production and are hence not suitable metabolic engineering targets.

Although targets are automatically filtered based on the overlap between flux distributions in different scenarios, the mean flux, and the range of the distribution during production, visual examination of distribution graphs is recommended as the final filter before experimental implementation.

#### 3.1.2 Effect of sampling parameters on target identification

Samples are taken in three scenarios: growth, slow growth and production. The user might choose an optimality constraint that determines the minimum growth or production in the growth and production scenarios respectively. The optimality parameter can take values from 0 to 1, where 0 indicates that flux distributions resulting in zero growth or production are allowed, and 1 indicates that only flux distributions with maximum growth or production are allowed. Increasing the optimality parameter tightens the flux constraints, reducing the feasible solution space and decreasing the number of over-expression targets (Figure 2). When the optimality parameter is increased, the minimum allowed production increases which results in an increased number of down-regulation targets (Figure 2). Reducing the solution space with the optimality parameter ensures that only relevant phenotypes are captured and a value of 0.9 is recommended as default. This optimality value ensures high production while allowing the sampling of sub-optimal phenotypes which increases the robustness of the predictions.

**Figure 2.**
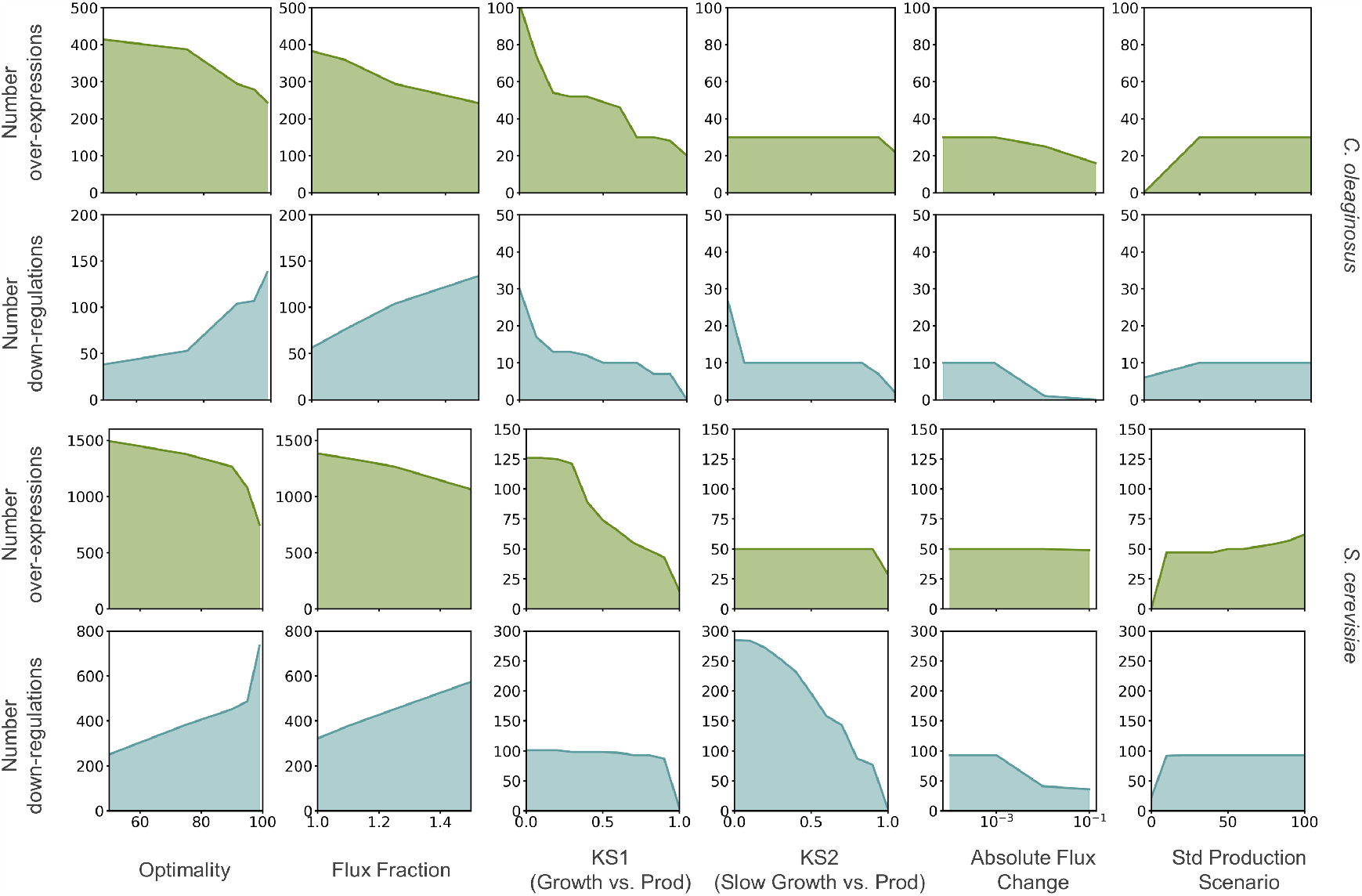
Effect of sampling and filtering parameters on the number of reported targets. The effect of sampling parameters was assessed using loose values for the filtering parameters (KS1 = KS2 ≤ 1, absolute flux change ≥0.001. std production ≤ 100). The effect of the filtering parameters was assessed on samples taken using optimality = 90% and flux fraction = 1.25, filtering parameters not under study were set to KS1 = KS2 ≤ 1, absolute flux change ≥ 0.001. std production ≤ 100.

Similarly, a flux faction parameter is used to limit the total sum of fluxes in the production scenario based on the total sum of fluxes in the growth scenario. This parameter can take values equal of bigger than 1, so higher flux fractions increase the available total flux and, therefore, enlarge the solution space. Lastiri-Pancardo et al. show that the fraction of unused proteome available for the expression of heterologous pathways is limited and influences production [25]. However, estimating this fraction is difficult and depends on the growth conditions [26]. Reducing the solution space with the flux fraction parameter as a proxi for proteome constraints reduces the risk of identifying unrealistic loops or excessively long pathways as engineering targets. We show that increasing the flux fraction parameter widens the solution space, increasing the number of down-regulation targets. At the same time, increasing the solution space might results in broader distributions that reduces mean flux differences between growth and production scenarios, reducing the number of possible over-expressions (Figure 2). We tested flux fraction values in the 1 to 1.5 range and found 1.25 to be a suitable value for the case studies here presented, although it can be adapted for other applications. Changing these parameters did not affect the sampling run time which is determined by the model size (94*±*1 min for *S. cerevisiae* model and 46*±*1 min for *C. oleaginosus* model on an Intel(R) Xeon(R) CPU E5-2650 v4 @2.2GHz).

#### 3.1.3 Effect of filtering parameters on target identification

Samples are evaluated based on the KS test, the mean flux in the different conditions and the variability of the mean flux in the production conditions (its standard deviation).

KS-test identifies whether samples belong to the same distributions (p-value) and the overlap between distributions (KS-statistic). Due to the large number of samples, the test is overpowered and p-values do not constitute a good filtering criteria even if corrections for multiple testing such as Bonferroni are applied. The KS-statistics determines the overlap between distributions, where a KS value of zero indicates complete overlap. This parameter largely affects the number of selected targets. KS1 indicates the overlap between the growth and production scenarios and ensures significantly different flux distributions among these conditions. Therefore, increasing KS1 reduces the number of over-expression and down-regulation targets (Figure 2). KS2 indicates the overlap between the slow growth and production scenarios. This condition is used to distinguish reactions in which flux decreases due to the decreased growth in the production scenario instead of the increased production. Therefore, increasing KS2 reduces the number of down-regulation targets. As default, the use of 0.75 for KS1 and KS2 is recommended but this value can be tuned based on the distribution of KS values obtained (see example in [GitLab](https://gitlab.com/wurssb/Modelling/sampling-tools/-/blob/master/example_filtering_yeast.ipynb?ref_type=heads)).

The absolute flux change between growth and production scenarios is used to rank the targets and can be used as an additional filter (Figure 2). Targeting reactions with considerable flux changes when growth or production are maximized, increases the chance of significant phenotype changes *in vivo*. Contrarily, low flow changes are unlikely to be achieved adjusting gene expression. A value of 0.001 mmol/gDW/h is used as default. Although we recommend the use of mean absolute change to favour reactions with larger fluxes, CFSA allows alternative filtering based on mean fold changes and mean relative changes of reaction fluxes.

The standard deviation of the flux distributions in the production scenario is used as an additional filter, since reactions with broad distributions are not considered relevant targets. Higher standard deviations indicate that production is not affected by the flux of the studied reaction. Decreasing the maximum allowed standard deviation therefore reduces the number of suggested targets for up and down-regulation. Considering that reactions with high mean fluxes also have a higher standard deviation, the default value of this parameter is set to 50.

### 3.2 Case studies

The number of possible metabolic-engineering targets obtained using CFSA for lipid production in *C. oleaginosus* and naringenin production in *S. cerevisiae* is presented in Table 2. This values were obtained using optimality = 0.90, flux fraction = 1.25, KS1 = KS2*≥*0.75, mean absolute change *≥*0.01 and standard deviation in production *≤*50. The complete list of target reactions can be found in [GitLab](https://gitlab.com/wurssb/Modelling/sampling-tools/-/blob/master/example_filtering_yeast.ipynb?ref_type=heads) (filtered_results.xlsx). The sections below elaborate on some of the targets found.

**Table 2:**
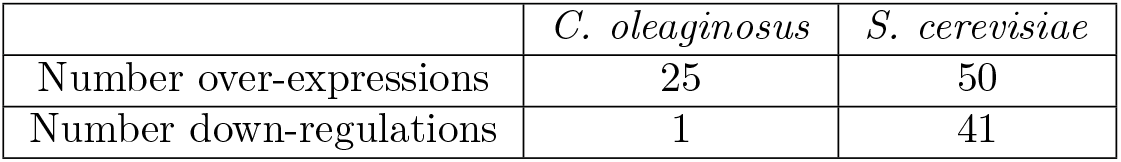
Number of targets obtained in the *C. oleaginosus* and *S. cerevisiae* case studies.

#### 3.2.1 Production of lipids by *C. oleaginosus*

*C. oleaginosus* is an oleaginous yeast able to accumulate lipids above 40% (w/w) of its biomass when growing on a nitrogen-limited cultivation medium [27]. The composition of the produced fatty acids, stored as triacylglycerols (TAGs) within lipid bodies in the cell, is comparable to commonly used plant-derived oils. Therefore it has been flagged as an auspicious MCF for sustainable lipid production at an industrial scale [28, 29]. The lipid accumulation initiates with the transport of citrate from the mitochondria to the cytosol where it is cleaved into acetyl-CoA by ATP citrate lyase (ACL). Acetyl-CoA is converted into malonyl-CoA by acetyl-CoA carboxylase (ACC) leading to the activation of the lipid synthesis and elongation pathways (Fig 3 A) [30, 31].

**Figure 3.**
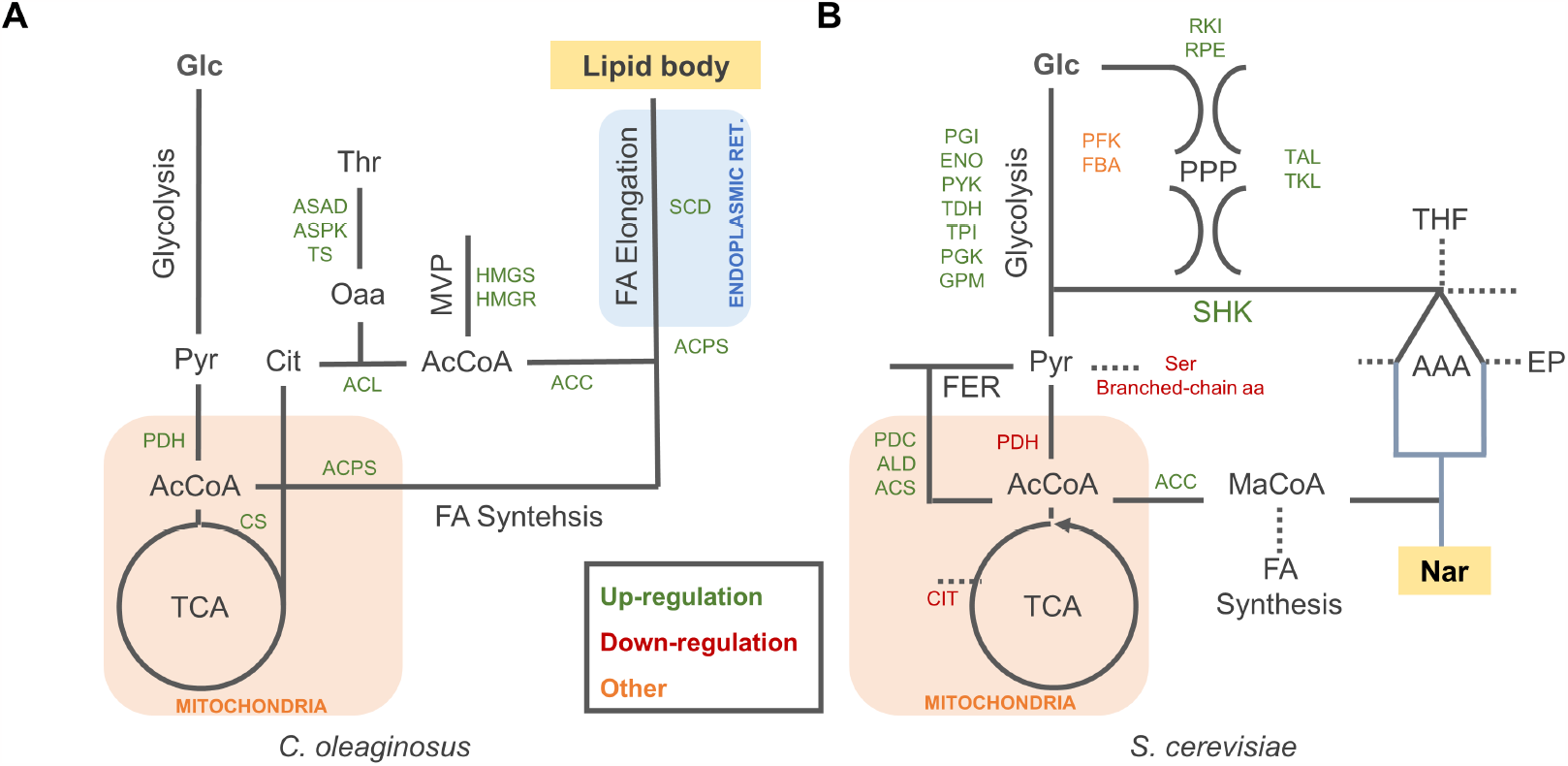
Summary of selected metabolic engineering targets for lipid production in *C. oleaginosus* (A) and naringenin (NAR) production in *S. cerevisiae* (B). Endogenous and heterologous metabolic pathways are simplified and presented in grey and blue respectively: TCA, tricarboxylic acid cycle; MVP, mevalonate pathway; FA, fatty acids; PPP, pentose phosphate pathway; SHK, shikimate pathway; FER, fermentation pathway; AAA, aromatic amino acid pathway; EP, Ehrlich pathway. Metabolite abbreviations: Glc, glucose; Py, pyruvate; AcCoA, acetyl-coenzyme A; Cit, Citrate; MaCoA, malonyl-coenzyme A.

**Figure 4.**
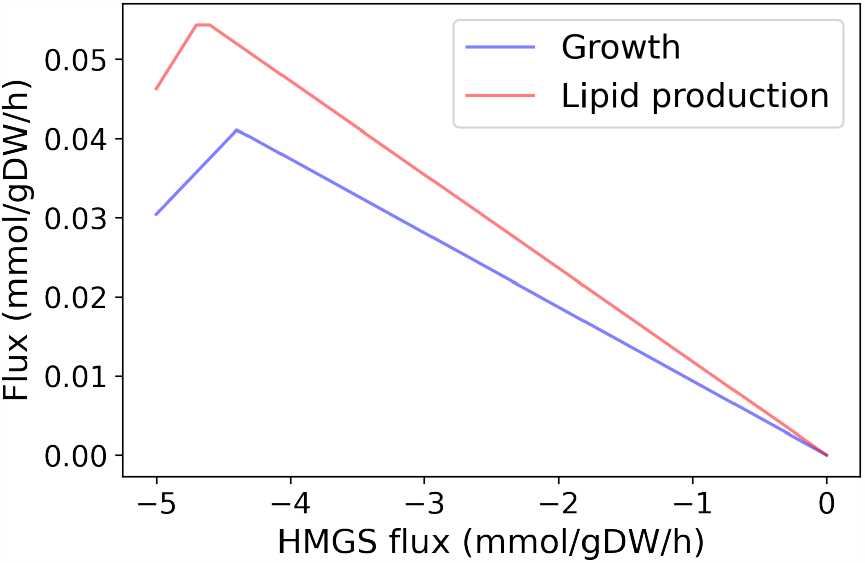
Effect of HMGS flux on growth and production fluxes. Fluxes were obtained using flux balance analysis fixing the bounds of the HMGS reaction (r_0599) and setting the growth (Biomass_nitrogen_depletion) or lipid production (Ex_lipid_body_cytosol) reactions as objective to maximize.

CFSA was applied to investigate metabolic engineering strategies for enhanced lipid production in *C. oleaginosus*. As a result, we obtained 1 candidate reaction for down-regulation and 25 candidate reactions (belonging to 11 groups) for over-expression. The complete list of reactions is available in [GitLab](https://gitlab.com/wurssb/Modelling/sampling-tools/-/blob/master/example_filtering_yeast.ipynb?ref_type=heads) (filtered_results.xlsx). The only down-regulation target found corresponds to ATP diphosphohydrolase that catalyzes the conversion of ATP to AMP and reflects the high energy requirements of lipid production compared to growth.

As expected, reactions from the fatty acid synthesis pathway (fatty acyl-acp synthase (ACPS) and stearoyl-CoA desaturase (SCD)) were suggested as over-expression targets (Figure 3A). Besides, key reactions ACL and ACC, which over-expression has improved lipid accumulation in other oleaginous yeast, were also predicted as targets [32, 33]. Additionally, pyruvate dehydrogenase (PDH), which connects the glycolytic pathway to the TCA cycle, and citrate synthase (CS), which synthesizes citrate from acetyl-CoA and oxaloacetate, were predicted as up-regulation targets to improve cytoplasmic citrate supply for lipid synthesis. However, CFSA did not predict down-regulation targets from the *β*-oxidation pathway that are commonly used to increase the availability of acetyl-CoA for fatty acid synthesis [34, 35].

Along with these experimentally validated targets, unanticipated reactions from the mevalonate pathway e.g., hydroxymethylglutaryl CoA synthase (HMGS) were suggested as beneficial up-regulations by CFSA. Although these reactions consume acetyl-CoA, simulations that reduce HMGS flux result in a diminished lipid production (Sup Figure 4). The complex relationship between the mevalonate pathway and lipid synthesis, intrinsic to the GEM used, is captured by CFSA as a possible strategy to increase fatty acid production.

Finally, CFSA suggested over-expression targets from amino acid metabolism, including reactions involved in threonine synthesis (aspartate-semialdehyde dehydrogenase (ASAD), aspartate kinase (ASPK), threonine synthase (TS)), as well as glutamate and glutamine synthases. Threonine synthesis requires cytosolic oxaloacetate which is produced during citrate conversion to acetyl-CoA by ACL. Therefore, the over-expression of the theronine synthesis pathway could improve lipid production balancing the over-production of oxaloacetate. Furthermore, Kim et al. suggested that upregulation of threonine synthesis could potentially increase the fluxes through the TCA cycle which increases citrate supply for acetyl-CoA synthesis [36].

#### 3.2.2 Production of naringenin by *S. cerevisiae*

*S. cerevisiae* is a model organism with in-depth genetic and physiological characterization, ample application in industrial bioprocesses and *Generally Regarded as Safe* (GRAS) status [37, 38]. Flavonoids such as naringenin are precursors of anthocyanins and have traditionally been used for fragrance, flavour, and colour in various food types [39, 40]. Naringenin is derived from the shikimate (SHK) pathway for aromatic amino acid biosynthesis, which starts with the condensation of erythrose-4-phosphate (E4P) and phosphoenolpyruvate (PEP) (Figure 3 B). Production of this compound requires the heterologous expression of 4CL, CHS, CHI, and either PAL, C4H and CPR for production from phenylalanine, or TAL for production from tyrosine (Table 1). Moreover, naringein production requires an appropriate supply of malonyl-CoA [41].

We used CFSA to find metabolic engineering strategies that improve naringenin production. We obtained 41 reaction candidates for down-regulation (belonging to 28 groups) and 50 targets for up-regulation (belonging to 35 groups). The complete list of reactions is available in [GitLab](https://gitlab.com/wurssb/Modelling/sampling-tools/-/blob/master/example_filtering_yeast.ipynb?ref_type=heads) (filtered_results.xlsx). Two proteomic datasets covering 48.4% of the genes and 23.6% of the reactions in the model were used as additional filtering step to prioritize the 34 detected proteins as down-regulation targets.

As expected, reactions from the shikimate and naringenin production pathways were predicted as over-expression targets, with a preference for the tyrosine branch. Reactions belonging to glycolysis and non-oxidative pentose phosphate pathway were also predicted as suitable targets (Figure 3B). Interestingly, CFSA suggested a decreased flux through the phosphofructokinase (PFK) and aldolase (FBA) reactions that result in the conversion of fructose-6-phosphate (F6P) to dihydroxyacetone-phosphate (DAP) and glyceraldehyde-3-phosphate (GAP). Instead, it proposed F6P conversion to E4P and DAP via sedoheptulose-1,7-biphosphate (S1,7bP). In this way E4P and DAP, that can be later converted to PEP, are simultaneously produced, feeding the shikimate pathway with its two precursors. In the cells, PFK and FBA are responsible for both pathways and conversion of F6P to DAP, and GAP is favoured due to a higher affinity of FBA towards F1,6-bP. According to the simulations, expressing *pfk* genes with higher affinity towards S7P such as *ppi-pfk* from *Clostridium thermosuccionogenes* [42] is suggested as a novel strategy to improve naringenin production.

CFSA also suggested experimentally validated strategies to increase the production of acetyl-CoA and malonyl-CoA syntheses such as down-regulation of PDH and CIT2 and up-regulation of PDCS, ALD, ACS and ACC (Fig 3B) [43]. However, it fails to predict the down-regulation of fatty acid synthesis as a strategy to improve malonyl-CoA availability [38, 43, 44]. Similarly, CFSA does not suggest the deletion of Ehrlich pathway (EP) genes, involved in the degradation of intermediates that have shown to improve the production of other aromatic compounds [40, 44–46].

The model suggests down-regulation of reactions involved in serine and branched chain amino acid synthesis (leucine, isoleucine and valine), likely to decrease the conversion of pyruvate into biomass components. Besides, down-regulation of reactions involving tetrahydrofolic acid (THF) and glycine are also suggested, probably to reduce chorismate consumption for THF formation (Fig 3B).

Last, CFSA suggested the over-expression of adenylate kinase (ADK) which has been reported to increase the production of malonyl-CoA derived products [47]. Similarly, although experimentally unrealistic, CFSA suggested down-regulation of ATP synthase and reactions from the electron transport chain, reflecting the lower energy requirements of production compared to growth.

## 4 Discussion

Inspired by other strain design algorithms that suggest metabolic engineering targets based on the comparison of predicted fluxes between *wild type* and production phenotypes [7–10], we present CFSA, a tool based on flux sampling that, by analyzing the complete GEM solution space, can guide the design of MCF. While current tools focus on design approaches for growth-couple production, we present a first tool that allows the design of growth-uncoupled production strategies. Besides, as opposed to previous tools, CFSA designs are based on the complete exploration of the solution space achieved using flux sampling [13] instead of non-unique FBA solutions, which ensures the full inspection of cell metabolism.

We apply CFSA to improve the production of lipids by *C. oleaginosus*, an endogenous product in an emerging MCF, as well as the production of naringenin in *S. cerevisiae*, a heterologous product in an established MCF. In both cases, CFSA suggested experimentally validated targets including evident up-regulations belonging to the product synthesis pathway and distant targets that improve precursor availability. It also suggested new engineering strategies involving alternative PPP reactions for naringenin synthesis and threonine and mevalonate pathways gene over-expressions for lipid production. Although the lack of mechanistic understanding of some of the suggested targets could question their implementation, it also highlights the potential of model-driven approaches to find non-obvious engineering strategies. Regarding down-regulations, CFSA fails to predict reported successful targets from pathways involved in the degradation of production pathway intermediates (e.g. Ehrlich pathway genes deletion for naringenin production [40, 44–46] or *β*-oxidation gene knock-outs for improved lipid production [34, 35]). This pitfall is shared with other GEM-based strain design approaches since, although active *in vivo*, these reactions remain inactive in GEM simulations.

In addition to the simulation of growth and production phenotypes, we include the simulation of slow growth. This scenario is used to differentiate between fluxes that change due to the simulated low-growth in the production scenario, from fluxes that are potentially related to increased production (Figure 1). The simulation of this phenotype is used as negative control reducing false positive target predictions (Figure 2). Other tools use FBA to maximize growth or production and use the obtained solution to compare reaction fluxes and identify engineering targets. Instead, we include the optimality parameter to ensure that not only the optimal flux profile is sampled, but less efficient behaviour is also tolerated leading to increased robustness.

CFSA is easy to implement and the filtering criteria used will result in a reduced list of potential targets for up-, down-regulations, and knock-outs for subsequent inspection. Default parameters can easily be modified according to user needs and additional filtering criteria such as the use of proteomic data can be integrated into the workflow. Other strain design methods provide minimal intervention strategies that, although useful in theory, might not return the expected results in practice. For example, when entire pathways are predicted as up-regulation targets, GEM cannot identify limiting reactions as enzyme kinetics and regulation are not considered in constraint-based models [4]. Instead, we provide a complete list of possible interventions and endow the user with additional information to make a decision based on the most feasible suggestions.

Disadvantages of CFSA include the difficulty of estimating the quantitative effect of the suggested manipulations, which are identified based on statistical testing. There is no guarantee that the effect size in flux correlates to the strength of the knock-down or over-expression required. Besides, targets are suggested as individual interventions and the effect of possible combinations of targets is not provided. Still, after a first round of CFSA, the algorithm can be re-applied using GEMs including desired interventions (i.e. with modified reaction bounds) to find potential complementary targets. As with other methods based on GEMs, errors in the models can lead to unexpected outcomes.

For example, loops in the metabolic network can lead to artificially high fluxes and thus false predictions. Often parsimonious flux balance analysis [17] or loop-less flux balance analysis [48, 49] is used to limit these errors, however, implementing these in flux sampling is non-trivial. We mimicked the parsimonious approach by adding a flux fraction parameter that constrains the total sum of fluxes in the production scenario based on the growing phenotype. However, this approach, while biologically reasonable for the reference condition at a high growth rate, does not necessarily apply to the production condition, as the assumption of minimal flux and thus protein usage does not hold for this artificial scenario. Instead, loop-less flux sampling approaches such as the loopless Artificially Centered Hit-and-Run on a Box algorithm (ll-ACHRB) [50] could be used to improve this process.

CFSA is the first strain design algorithm based on flux sampling that explores the whole solution space of a GEM and suggests metabolic engineering targets for growth-uncoupled production. Its robustness, simplicity, and flexibility make it ideal to complement and systematize MCF design.

CFSA predictions, including non-obvious targets, can be sequentially tested using high-throughput approaches such as automated platforms and biosensor-aid screening accelerating and broadening the strain design process.

## 5 Acknowledgement

This project has been funded by the Netherlands Organization for Scientific Research (NWO; project number GSGT.2019.008) and the Dutch Ministry of Agriculture through the “TKI-toeslag” project LWV19221 “Tailor-made microbial oils and fatty acids”.

## 6 Supplementary Figures

## References

[1] S. Y. Lee et al. “A comprehensive metabolic map for production of bio-based chemicals”. en. In: Nature Catalysis 2.1 (2019). Number: 1 Publisher: Nature Publishing Group, pp. 18–33. doi: 10.1038/s41929-018-0212-4. url: https://www.nature.com/articles/s41929-018-0212-4.

[2] J. S. Cho et al. “Designing Microbial Cell Factories for the Production of Chemicals”. In: JACS Au 2.8 (2022). Publisher: American Chemical Society, pp. 1781–1799. doi: 10.1021/jacsau.2c00344. url: 10.1021/jacsau.2c00344.

[3] X. Fang, C. J. Lloyd, and B. O. Palsson. “Reconstructing organisms in silico: genome-scale models and their emerging applications”. en. In: Nature Reviews Microbiology 18.12 (2020). Number: 12 Publisher: Nature Publishing Group, pp. 731–743. doi: 10.1038/s41579-020-00440-4. url: https://www.nature.com/articles/s41579-020-00440-4.

[4] D. Machado and M. J. Herrgard. “Co-Evolution of Strain Design Methods Based on Flux Balance and Elementary Mode Analysis”. In: Metabolic Engineering Communications 2 (2015), pp. 85–92. doi: 10.1016/j.meteno.2015.04.001.

[5] C. T. Trinh, A. Wlaschin, and F. Srienc. “Elementary Mode Analysis: A Useful Metabolic Pathway Analysis Tool for Characterizing Cellular Metabolism”. In: Applied Microbiology and Biotechnology 81.5 (2009), pp. 813–826. doi: 10.1007/s00253-008-1770-1.

[6] A. von Kamp and S. Klamt. “Enumeration of Smallest Intervention Strategies in Genome-Scale Metabolic Networks”. In: PLOS Computational Biology 10.1 (2014), e1003378. doi: 10.1371/journal.pcbi.1003378.

[7] A. P. Burgard, P. Pharkya, and C. D. Maranas. “Optknock: A Bilevel Programming Framework for Identifying Gene Knockout Strategies for Microbial Strain Optimization”. In: Biotechnology and Bioengineering 84.6 (2003), pp. 647–657. doi: 10.1002/bit.10803.

[8] N. Tepper and T. Shlomi. “Predicting Metabolic Engineering Knockout Strategies for Chemical Production: Accounting for Competing Pathways”. In: Bioinformatics 26.4 (2010), pp. 536–543. doi: 10.1093/bioinformatics/btp704.

[9] K. R. Patil et al. “Evolutionary Programming as a Platform for in Silico Metabolic Engineering”. In: BMC Bioinformatics 6.1 (2005), pp. 1–12. doi: 10.1186/1471-2105-6-308.

[10] K. Jensen et al. “OptCouple: Joint Simulation of Gene Knockouts, Insertions and Medium Modifications for Prediction of Growth-Coupled Strain Designs”. In: Metabolic Engineering Communications 8 (2019), e00087. doi: 10.1016/j.mec.2019.e00087.

[11] S. Ranganathan, P. F. Suthers, and C. D. Maranas. “OptForce: An Optimization Procedure for Identifying All Genetic Manipulations Leading to Targeted Overproductions”. In: PLOS Computational Biology 6.4 (2010), e1000744. doi: 10.1371/journal.pcbi.1000744.

[12] S. Jiang et al. “OptDesign: Identifying Optimum Design Strategies in Strain Engineering for Biochemical Production”. In: ACS Synthetic Biology (2022), acssynbio.1c00610. doi: 10.1021/ACSSYNBIO.1C00610. url: https://pubs.acs.org/doi/full/10.1021/acssynbio.1c00610.

[13] H. A. Herrmann et al. “Flux sampling is a powerful tool to study metabolism under changing environmental conditions”. In: npj Systems Biology and Applications 5.1 (2019), pp. 1–8. doi: 10.1038/s41540-019-0109-0.

[14] S. Ravi and R. Gunawan. “?FBA—Predicting metabolic flux alterations using genome-scale metabolic models and differential transcriptomic data”. In: PLOS Computational Biology 17.11 (2021), e1009589. doi: 10.1371/JOURNAL.PCBI.1009589. url: https://journals.plos.org/ploscompbiol/article?id=10.1371/journal.pcbi.1009589.

[15] T. M. Lo et al. “A Two-Layer Gene Circuit for Decoupling Cell Growth from Metabolite Production”. In: Cell Systems 3.2 (2016), pp. 133–143. doi: 10.1016/j.cels.2016.07.012. url: 10.1016/j.cels.2016.07.012.

[16] K. Raj, N. Venayak, and R. Mahadevan. “Novel two-stage processes for optimal chemical production in microbes”. In: Metabolic Engineering 62 (2020), pp. 186–197. doi: 10.1016/j.ymben.2020.08.006. url: https://www.sciencedirect.com/science/article/pii/S1096717620301269.

[17] N. E. Lewis et al. “Omic Data from Evolved E. Coli Are Consistent with Computed Optimal Growth from Genome-Scale Models”. In: Molecular Systems Biology 6.1 (2010), p. 390. doi: 10.1038/msb.2010.47.

[18] W. Megchelenbrink, M. Huynen, and E. Marchiori. “optGpSampler: An Improved Tool for Uniformly Sampling the Solution-Space of Genome-Scale Metabolic Networks”. In: PLoS ONE 9.2 (2014). Ed. by S. Rogers, e86587. doi: 10.1371/journal.pone.0086587.

[19] A. Ebrahim et al. “COBRApy: COnstraints-Based Reconstruction and Analysis for Python.” In: BMC systems biology 7.1 (2013), p. 74. doi: 10.1186/1752-0509-7-74.

[20] M. K. Cowles and B. P. Carlin. “Markov Chain Monte Carlo Convergence Diagnostics: A Comparative Review”. In: Journal of the American Statistical Association 91.434 (1996), pp. 883–904. doi: 10.1080/01621459.1996.10476956.

[21] N. Pham et al. “Genome-Scale Metabolic Modeling Underscores the Potential of Cutaneotrichosporon Oleaginosus ATCC 20509 as a Cell Factory for Biofuel Production”. In: Biotechnology for Biofuels 14.1 (2021), p. 2. doi: 10.1186/s13068-020-01838-1.

[22] H. Lu et al. “A Consensus S. Cerevisiae Metabolic Model Yeast8 and Its Ecosystem for Comprehensively Probing Cellular Metabolism”. In: Nature Communications 10.1 (2019), pp. 1–13. doi: 10.1038/s41467-019-11581-3.

[23] R. Costenoble et al. “Comprehensive Quantitative Analysis of Central Carbon and Amino-acid Metabolism in Saccharomyces Cerevisiae under Multiple Conditions by Targeted Proteomics”. In: Molecular Systems Biology 7.1 (2011), p. 464. doi: 10.1038/msb.2010.122.

[24] P. .-. Lahtvee et al. “Absolute Quantification of Protein and mRNA Abundances Demonstrate Variability in Gene-Specific Translation Efficiency in Yeast”. In: Cell Systems 4.5 (2017), 495–504.e5. doi: 10.1016/j.cels.2017.03.003.

[25] G. Lastiri-Pancardo et al. “A quantitative method for proteome reallocation using minimal regulatory interventions”. In: Nature Chemical Biology 16.9 (2020), pp. 1026–1033.

[26] I. E. Elsemman et al. “Whole-cell modeling in yeast predicts compartment-specific proteome constraints that drive metabolic strategies”. In: Nature Communications 2022 13:1 13.1 (2022), pp. 1–12. doi: 10.1038/s41467-022-28467-6. url: https://www.nature.com/articles/s41467-022-28467-6.

[27] Z. E. Duman-Ozdamar et al. “Tailoring and optimizing fatty acid production by oleaginous yeasts through the systematic exploration of their physiological fitness”. In: Microbial Cell Factories 21.1 (2022), pp. 1–13. doi: 10.1186/S12934-022-01956-5/FIGURES/3. url: https://microbialcellfactories.biomedcentral.com/articles/10.1186/s12934-022-01956-5.

[28] J. Zhang et al. “Microbial Lipid Production by the Oleaginous Yeast Cryptococcus Curvatus O3 Grown in Fed-Batch Culture”. In: Biomass and Bioenergy 35.5 (2011), pp. 1906–1911. doi: 10.1016/j.biombioe.2011.01.024.

[29] N. Di Fidio et al. “Cutaneotrichosporon Oleaginosus: A Versatile Whole-Cell Biocatalyst for the Production of Single-Cell Oil from Agro-Industrial Wastes”. In: Catalysts 11.11 (2021), p. 1291. doi: 10.3390/catal11111291.

[30] F. Bracharz et al. “Opportunities and Challenges in the Development of Cutaneotrichosporon Oleaginosus ATCC 20509 as a New Cell Factory for Custom Tailored Microbial Oils”. In: Microbial Cell Factories 16.1 (2017), p. 178. doi: 10.1186/s12934-017-0791-9.

[31] C. Gorner et al. “Genetic Engineering and Production of Modified Fatty Acids by the Non-Conventional Oleaginous Yeast Trichosporon Oleaginosus ATCC 20509”. In: Green Chemistry 18.7 (2016), pp. 2037–2046. doi: 10.1039/C5GC01767J.

[32] H. Zhang et al. “Enhanced Lipid Accumulation in the Yeast Yarrowia Lipolytica by Over-Expression of ATP:Citrate Lyase from Mus Musculus”. In: Journal of Biotechnology 192 (2014), pp. 78–84. doi: 10.1016/j.jbiotec.2014.10.004.

[33] F. X. Yan et al. “Overexpression of Δ12, Δ15-Desaturases for Enhanced Lipids Synthesis in Yarrowia Lipolytica”. In: Frontiers in Microbiology 11 (2020). doi: 10.3389/fmicb.2020.00289.

[34] C. Madzak. “Yarrowia lipolytica Strains and Their Biotechnological Applications: How Natural Biodiversity and Metabolic Engineering Could Contribute to Cell Factories Improvement”. In: Journal of Fungi 7.7 (2021). doi: 10.3390/jof7070548. url: https://www.mdpi.com/2309-608X/7/7/548.

[35] J. Blazeck et al. “Heterologous production of pentane in the oleaginous yeast Yarrowia lipolytica”. In: Journal of Biotechnology 165.3 (2013), pp. 184–194. doi: 10.1016/j.jbiotec.2013.04.003. url: https://www.sciencedirect.com/science/article/pii/S0168165613001831.

[36] M. Kim et al. “In Silico Identification of Metabolic Engineering Strategies for Improved Lipid Production in Yarrowia Lipolytica by Genome-Scale Metabolic Modeling”. In: Biotechnology for Biofuels 12.1 (2019), pp. 1–14. doi: 10.1186/s13068-019-1518-4.

[37] M. Gottardi et al. “Pathway Engineering for the Production of Heterologous Aromatic Chemicals and Their Derivatives in Saccharomyces Cerevisiae: Bioconversion from Glucose”. In: FEMS Yeast Research 17.4 (2017), p. 35. doi: 10.1093/femsyr/fox035.

[38] J. Lian, S. Mishra, and H. Zhao. “Recent Advances in Metabolic Engineering of Saccharomyces Cerevisiae: New Tools and Their Applications”. In: Metabolic Engineering 50 (2018), pp. 85–108. doi: 10.1016/J.YMBEN.2018.04.011.

[39] P. S. Nigam and J. S. Luke. “Food Additives: Production of Microbial Pigments and Their Antioxidant Properties”. In: Current Opinion in Food Science 7 (2016), pp. 93–100. doi: 10.1016/J.COFS.2016.02.004.

[40] A. Rodriguez et al. “Metabolic Engineering of Yeast for Fermentative Production of Flavonoids”. In: Bioresource Technology 245 (2017), pp. 1645–1654. doi: 10.1016/j.biortech.2017.06.043.

[41] X. Lyu et al. “Enhancement of Naringenin Biosynthesis from Tyrosine by Metabolic Engineering of Saccharomyces Cerevisiae”. In: Journal of Agricultural and Food Chemistry 65.31 (2017), pp. 6638–6646. doi: 10.1021/acs.jafc.7b02507.

[42] J. G. G. Koendjbiharie et al. “The Pentose Phosphate Pathway of Cellulolytic Clostridia Relies on 6-Phosphofructokinase Instead of Transaldolase”. In: Journal of Biological Chemistry 295.7 (2020), pp. 1867–1878. doi: 10.1074/jbc.RA119.011239.

[43] Y. Chen et al. “Coupled Incremental Precursor and Co-Factor Supply Improves 3-Hydroxypropionic Acid Production in Saccharomyces Cerevisiae”. In: Metabolic Engineering 22 (2014), pp. 104–109. doi: 10.1016/j.ymben.2014.01.005.

[44] S. Li et al. “Recent Progress in Metabolic Engineering of Saccharomyces Cerevisiae for the Production of Malonyl-CoA Derivatives”. In: Journal of Biotechnology 325 (2021), pp. 83–90. doi: 10.1016/j.jbiotec.2020.11.014.

[45] L. Milke et al. “Production of Plant-Derived Polyphenols in Microorganisms: Current State and Perspectives”. In: Applied Microbiology and Biotechnology 102.4 (2018), pp. 1575–1585. doi: 10.1007/s00253-018-8747-5.

[46] N. D. Gold et al. “Metabolic Engineering of a Tyrosine-Overproducing Yeast Platform Using Targeted Metabolomics”. In: Microbial Cell Factories 14 (2015), p. 73. doi: 10.1186/s12934-015-0252-2.

[47] R. Ferreira et al. “Model-Assisted Fine-Tuning of Central Carbon Metabolism in Yeast through dCas9-Based Regulation”. In: ACS Synthetic Biology 8.11 (2019), pp. 2457–2463. doi: 10.1021/acssynbio.9b00258.

[48] J. Schellenberger, N. E. Lewis, and B. Ø. Palsson. “Elimination of Thermodynamically Infeasible Loops in Steady-State Metabolic Models”. In: Biophysical Journal 100.3 (2011), pp. 544–553. doi: 10.1016/j.bpj.2010.12.3707.

[49] A. A. Desouki et al. “CycleFreeFlux: Efficient Removal of Thermodynamically Infeasible Loops from Flux Distributions”. In: Bioinformatics 31.13 (2015), pp. 2159–2165. doi: 10.1093/bioinformatics/btv096.

[50] P. A. Saa and L. K. Nielsen. “Ll-ACHRB: A Scalable Algorithm for Sampling the Feasible Solution Space of Metabolic Networks”. In: Bioinformatics 32.15 (2016), pp. 2330–2337. doi: 10.1093/bioinformatics/btw132.

